# Spatial organization of the transcriptome in individual neurons

**DOI:** 10.1101/2020.12.07.414060

**Authors:** Guiping Wang, Cheen-Euong Ang, Jean Fan, Andrew Wang, Jeffrey R. Moffitt, Xiaowei Zhuang

## Abstract

Neurons are highly polarized cells with complex neurite morphology. Spatial organization and local translation of RNAs in dendrites and axons play an important role in many neuronal functions. Here we performed super-resolution spatial profiling of RNAs inside individual neurons at the genome scale using multiplexed error-robust fluorescence *in situ* hybridization (MERFISH), and mapped the spatial organization of up to ∼4,200 RNA species (genes) across multiple length scales, ranging from sub-micrometer to millimeters. Our data generated a quantitative intra-neuronal atlas of RNAs with distinct transcriptome compositions in somata, dendrites, and axons, and revealed diverse sub-dendritic distribution patterns of RNAs. Moreover, our spatial analysis identified distinct groups of genes exhibiting specific spatial clustering of transcripts at the sub-micrometer scale that were dependent on protein synthesis and differentially dependent on synaptic activity. Overall, these data provide a rich resource for characterizing the subcellular organization of the transcriptome in neurons with high spatial resolution.

---

The spatial organization of RNAs inside cells is a powerful mechanism for post-transcriptional regulation (*1*–*6*). This is particularly important for neurons, which are highly polarized cells that extend long axons and dendrites from somata. These neurites respond to local signals with a high degree of autonomy. For example, axons can navigate correctly even after being severed from their originating somata, and likewise, isolated dendrites can respond to neurotrophic factors and undergo synaptic plasticity (*7, 8*). RNAs localized to neurites, which enable local production of proteins, form a basis for such autonomy and are critical to key neuronal functions such as synaptic plasticity, axon branching and pathfinding, and retrograde signaling (*1*–*3, 5, 6*). Recent evidence has shown that differential enrichment of RNAs in different sub-neuronal compartments is a key determinant of the proteome composition in neurites (*9, 10*). High-resolution single-molecule RNA imaging, albeit for a relatively small number of genes, has revealed different spatial distributions of RNAs inside neurons (*2, 6*). Changes in dendritic RNA distributions induced by removal of the localizing signal in the 3’UTR of RNAs or local reduction in mRNA half-life can substantially affect local protein distribution and functions (*10*– *12*). A systematic, quantitative and high-resolution characterization of the intracellular spatial organization of the transcriptome would thus be critical for our understandings of a variety of neuronal functions. Up to thousands of RNA species have been found present or enriched in neurites by microarray and sequencing analyses (*1, 5, 6*). However, due to the limited spatial resolution of these genomics approaches, quantitative characterization of the sub-neuronal spatial distributions, in particular at the sub-neurite level, is still lacking for most RNAs. Moreover, the spatial relationship between different RNA species remains largely unknown.

Here we used MERFISH, a single-cell transcriptome imaging method (*13*) that massively multiplexes single-molecule FISH (*14, 15*) using error-robust barcoding, combinatorial labeling and sequential imaging, to map the subcellular distributions of up to ∼4,200 RNA species (genes) in hundreds to thousands of individual neurons. Our spatially resolved single-cell transcriptomic analysis provides the copy numbers and spatial distributions of individual RNA species in somata, dendrites and axons. We observed that a large number of RNA species were enriched in dendrites, encoding proteins of diverse functions, ranging from dendritic transport to synaptic plasticity. In axons, several kinesin-related mRNAs were expressed at exceptionally high levels, up to tens of axonal copies per cell, whereas the vast majority of other genes were expressed in axons at very low copy numbers. At the sub-dendritic scale, we observed diverse spatial distribution patterns along the proximal-distal axis of dendrites for different groups of genes. Down to the sub-micrometer scale, we identified distinct clusters of genes exhibiting spatial clustering among their transcripts in a manner that were dependent on protein synthesis and differentially dependent on synaptic activity.

To map the spatial distributions of RNAs in neurons, we performed two sets of three-dimensional (3D) MERFISH imaging experiments, targeting 950 and 4,209 genes, respectively, in cultured mouse hippocampal neurons at 18 days in vitro (DIV) unless otherwise mentioned. In the first set of experiments, we selected 950 genes related to neuron projections, neurite development and synapses. We also included well-characterized cell type markers, such as *Slc17a7, Gad1*/*2, Gfap, Pdgfra*, to identify excitatory neurons, inhibitory neurons, astrocytes and oligodendrocytes, respectively. We imaged these genes using MERFISH with a 32-bit Hamming distance 4 (HD4), Hamming weight 4 (HW4) code capable of error correction (*13*). Of the 1,240 total possible HD4, HW4 barcodes, we randomly assigned 950 of them to the 950 targeted RNA species and left 290 barcodes as blank controls. Neurons have compact somata, resulting in a high density of RNA molecules inside somata. We thus used expansion microscopy (*16, 17*) with a 2-fold linear expansion ratio, to help resolve nearby RNA molecules (*18, 19*). We further immuno-labeled dendritic and axonal markers, MAP2 and Tau, respectively, to map RNAs to various compartments of neurons (Fig. 1A).

**Fig. 1.**
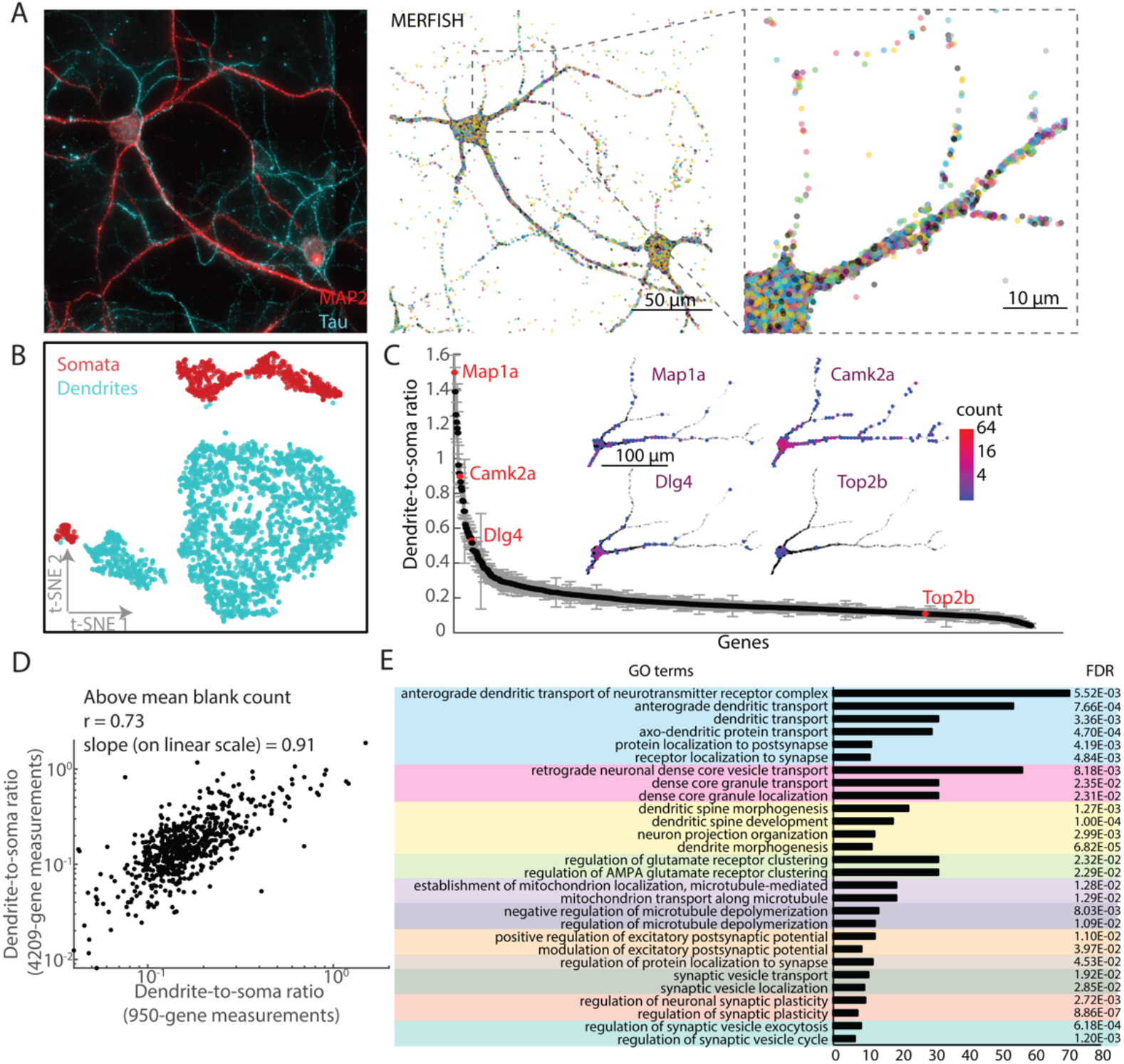
Identification and quantification of RNAs enriched in dendrites. (**A**) (Left) MAP2 and Tau immunofluorescence images from a sub-region of a 4 mm-by-4 mm imaged area showing two representative neurons. (Middle) MERFISH image of 950 genes in the same region. Each colored dot represents a single RNA molecule colored by its gene identity. Scale bar: 100 µm after expansion or 50 µm before expansion. (Right) Zoomed-in image of the region in the dash box from the middle panel. (**B**) T-distributed stochastic neighbor embedding (tSNE) for individual somatic (red) and dendritic (cyan) compartments of both excitatory and inhibitory neurons based on the transcriptome profiles. All annotated somatic and dendritic regions (617 somata and 2274 dendritic branches, from 4 biological replicates) are shown. (**C**) Dendrite-to-soma ratios for individual RNA species measured in all annotated neurons in the 950-gene measurements, with RNA species (genes) rank-ordered based on their dendrite-to-soma ratio. The error bar represents the SEM determined from 4 biological replicates. (inset) RNA spatial maps of four example RNA species highlighted in red in a single neuron. Each hexagon represents a 5 µm by 5 µm region with the color corresponding to the local copy number of the indicated RNA species at the location. The other RNA species are marked as semi-transparent black dots. Scale bar: 100 µm before expansion. (**D**) Scatter plot of dendrite-to-soma ratios of common RNAs measured in the 4,209-gene experiments versus those measured in the 950-gene experiments. The Pearson correlation coefficient (r) is 0.73 and the slope of linear regression of the data is 0.91 (both determined with data on the linear scale). (**E**) Representative groups of GO terms that are preferentially enriched among the RNA species substantially enriched in dendrites (i.e. RNA species with fold of enrichment in dendrites > 1.5 fold; p-value < 0.05), as determined from the 4209-gene measurements. GO term groups are color-coded, and their fold of enrichment and Benjamini-Hochberg False Discovery Rates (FDR) are shown. GO terms enriched among the genes substantially enriched in dendrites and GO terms enriched among the rest of the genes were separately determined, and representative GO terms that are either uniquely enriched among dendritically enriched genes or with fold of enrichment at least two-fold higher than those in the other group are shown.

Out of the 950 targeted RNA species, 90% (853 RNA species) had detected copy numbers per cell that were consistently above the blank control (i.e. the mean copy number of blank control barcodes not assigned to any RNA) in all four biological replicates. The average copy numbers of individual genes per cell were highly reproducible across biological replicates (Pearson correlation coefficient r = 0.94, Fig. S1). To assess how the high total RNA density of the 950 genes affected our detection efficiency, we performed lower multiplexity, 100-gene MERFISH measurements for comparison. Based on the relatively low density of RNA molecules in these lower multiplexity measurements (<0.5 spot per µm^2^) and our previous of MERFISH experiments with similar RNA density and similar gene number (*18, 20*), we anticipate that the detection efficiency of these 100-gene measurements to be ∼100%. Due to molecular crowding (*18*), the per-gene detection efficiency of the 950-gene measurements were 85% of that of 100-gene measurements for transcripts detected in neurites and 38% for transcripts detected in somata, determined based on the copy number per gene detected for the genes commonly present in both 100- and 950-gene libraries (Fig. S2).

To facilitate visual inspection of the MERFISH data, we developed a Javascript-based web application to enable interactive exploration of spatial expression patterns of genes of interest, which allows efficient rendering of millions of individual molecules in space and real-time highlighting of different RNA species in single cells across a wide range of spatial scales (from <100 nm to >1 mm) overlaid on immunofluorescence images of the same region (Fig. S3).

To facilitate quantitative analysis, we classified dendritic regions based on MAP2 immunofluorescence signals, and manually annotated the somatic and dendritic regions that can be traced to individual neurons. We detected a median of ∼11,000 RNA molecules per neuron (25-75% quantile range: 9,050-14,500) and ∼17% of them were in annotated dendrites. At the single-compartment level, out of the 853 RNA species whose detected copy numbers per cell were above the blank control, we detected a median of 740 RNA species in individual somata, and a median of 474 RNA species in the annotated dendritic regions of individual neurons.

Notably, dendrites exhibited distinct transcriptome compositions compared to somata (Fig. 1B and Fig. S4). To quantify this effect, we determined, for each RNA species, the ratio of the copy number in dendrites to that in somata (hereafter referred to as the dendrite-to-soma ratio), after taking into consideration the different detection efficiencies in dendrites and somata (Fig. S2) and the fraction of dendrites annotated (See Supplementary Materials). The dendrite-to-soma ratios spanned a wide range of values for different RNA species (Fig. 1C). This ratio was reproducible between biological replicates (Fig. S5A). The top 10% of RNA species with the highest dendrite-to-soma ratios includes the previously known “gold standard” dendritic RNAs such as *Camk2a, Shank1* and *Dlg4*, as well as other RNAs that span a wide range of functional categories, such as RNAs encoding post-synaptic density proteins (*Homer2, Dlg2*), calcium sensor and signaling molecules (*Ncdn, Ncs1, Nsmf*), kinases, kinase inhibitors and phosphatases (*Cdk16, Map2k2, Prkcz, Cit, Camk2n1, Ppp1r9b, Ppp2r5b, Ppp1r2*), ion channels (*Kcnab2, Hcn2, Asic2*), cytoskeletal proteins and regulators (*Nefl, Wasf3, Evl*), ribosomal proteins (*Rpsa, Rps3*), and motor proteins (*Trak1*/*2, Dynll1, Myo5a, Kif5a, Kif5b, Kif5c*), etc. We further compared soma-dendrite RNA distributions in excitatory neurons (as marked by the expression of *Slc17a7*, Fig. S4) and inhibitory neurons (as marked by expression of *Gad1* and/or *Gad2*, Fig. S4). Interestingly, despite the fact that many genes were differentially expressed in excitatory and inhibitory neurons, we observed highly correlated dendrite-to-soma ratios for the imaged RNA species between these different neuronal types (Fig. S5B, Pearson correlation coefficient r = 0.88).

Dendritic transcripts may be developmentally regulated (*21*). We investigated expression level changes of these 950 genes in the somata and dendrites of excitatory neurons upon neuronal maturation by performing MERFISH experiments on neurons at two different maturation stages (DIV 10 vs 18). Approximately equal numbers of genes were upregulated and downregulated in somata during maturation (Fig. S6A). In contrast, most genes were upregulated in dendrites upon neuronal maturation (Fig. S6B). The upregulated transcripts in dendrites included not only transcripts with high dendrite-to-soma ratios, but also many transcripts with medium to low dendrite-to-soma ratios (Fig. S6C).

To gain a more comprehensive view of dendritic enrichment of RNAs in dendrites of mature neurons, we next imaged 4,209 genes, including both cell type markers and genes contained in general neuron-related gene ontology (GO) terms, in DIV18 neurons. We performed MERFISH experiments of these genes using a 49-bit HD4, HW4 error-correcting code (4508 total possible barcodes: 4209 assigned to targeted genes and 299 assigned to blank controls). Of the 4,209 RNA species targeted, 86% (3,615 RNA species) had detected copy numbers per cell that were consistently above the mean copy number of the blank control barcodes in both of the biological replicates measured, and the average copy numbers of individual genes were reproducible between the two replicates (Fig. S7). Because of the substantially higher density of RNA molecules in this gene library, and hence the more severe overlap of signals among neighboring RNA molecules, in particular in the somatic region, the detected copy number per gene were reduced compared to the 950-gene measurements: for the RNA species that were included in both gene libraries, the copy numbers per gene detected in the 4,209-gene measurement were 65%, 42% and 15% of those detected in the 950-gene measurements for transcripts observed in axons, dendrites and somata, respectively (Fig. S8). Despite the lower detection efficiency, the copy numbers per gene detected in the 4,209-gene measurements were highly correlated with those measured in the 950-gene measurements for all three neuronal compartments (Fig. S8). In subsequent analysis, we considered the difference in detection efficiency in different neuronal compartments. The dendrite-to-soma ratios measured in the 4,209-gene measurements were consistent with those measured with the 950-gene measurements, with a Pearson correlation coefficient of 0.73 and a linear regression slope of 0.91 for genes with copy number above the mean blank control (Fig. 1D).

The large number of genes included in the 4,209-gene library allowed us to better estimate the baseline dendrite-to-soma ratio and determine which genes were enriched in dendrites. For genes annotated to be related to neuronal cell body and unrelated to neurites and synapses, the median dendrite-to-soma ratio was 0.16 (See Supplementary Materials). To determine the genes that were enriched in dendrites, we compared the dendritic RNA counts for each gene to a permutation control, where we randomly shuffled the gene identities of detected RNA molecules while preserving the total copy number of each gene. We then considered a gene enriched in dendrites if the measured dendritic count was consistently higher than 95% of the randomization controls (i.e. enrichment p-value < 0.05). Based on this criterion, ∼12% of the 3,615 RNA species above the mean black control were enriched in dendrites. We then performed GO term analysis for genes that were substantially enriched in dendrites (with >1.5-fold enrichment; p-value < 0.05) and compared with GO term analysis for the rest of the measured genes. We found that these substantially dendritically enriched genes were preferentially enriched for GO terms related to a variety of dendritic and synaptic functions, such as dendritic transport, protein localization to postsynapses, dendritic spine morphogenesis and development, regulation of glutamate receptor clustering, regulation of postsynaptic potential, synaptic vesicle transport and localization, regulation of synaptic plasticity, regulation of synaptic vesicle exocytosis and cycle, etc (Fig. 1E).

Next, we quantified the axonal RNA copy numbers of the 4,209 imaged genes, which include ∼2,000 genes previous identified to be present in axonal regions (*22*–*24*). We identified axonal regions by both Tau staining and separation from MAP2 and glia markers (*Gfap, Pdgfra*) (Fig. 2A). Although we observe many of these RNA species in axons, we found that most of these RNA species existed in very low copy number in axons (>95% and >70% of the 4209 genes had <1 and <0.1 axonal copies of RNAs per cell, respectively). To quantify the axonal enrichment for the imaged genes, we computed the enrichment p-values of individual RNA species in axons, and more specifically in axons versus dendrites, by comparing our measured results to randomization controls, similar to what was described above for dendritic enrichment characterization. We found that, out of 3,615 RNA species above black controls, 4.5% of the RNA species were enriched in axons and 4% of the RNA species were specifically enriched in axons versus dendrites (enrichment p-value <0.05).

**Fig. 2.**
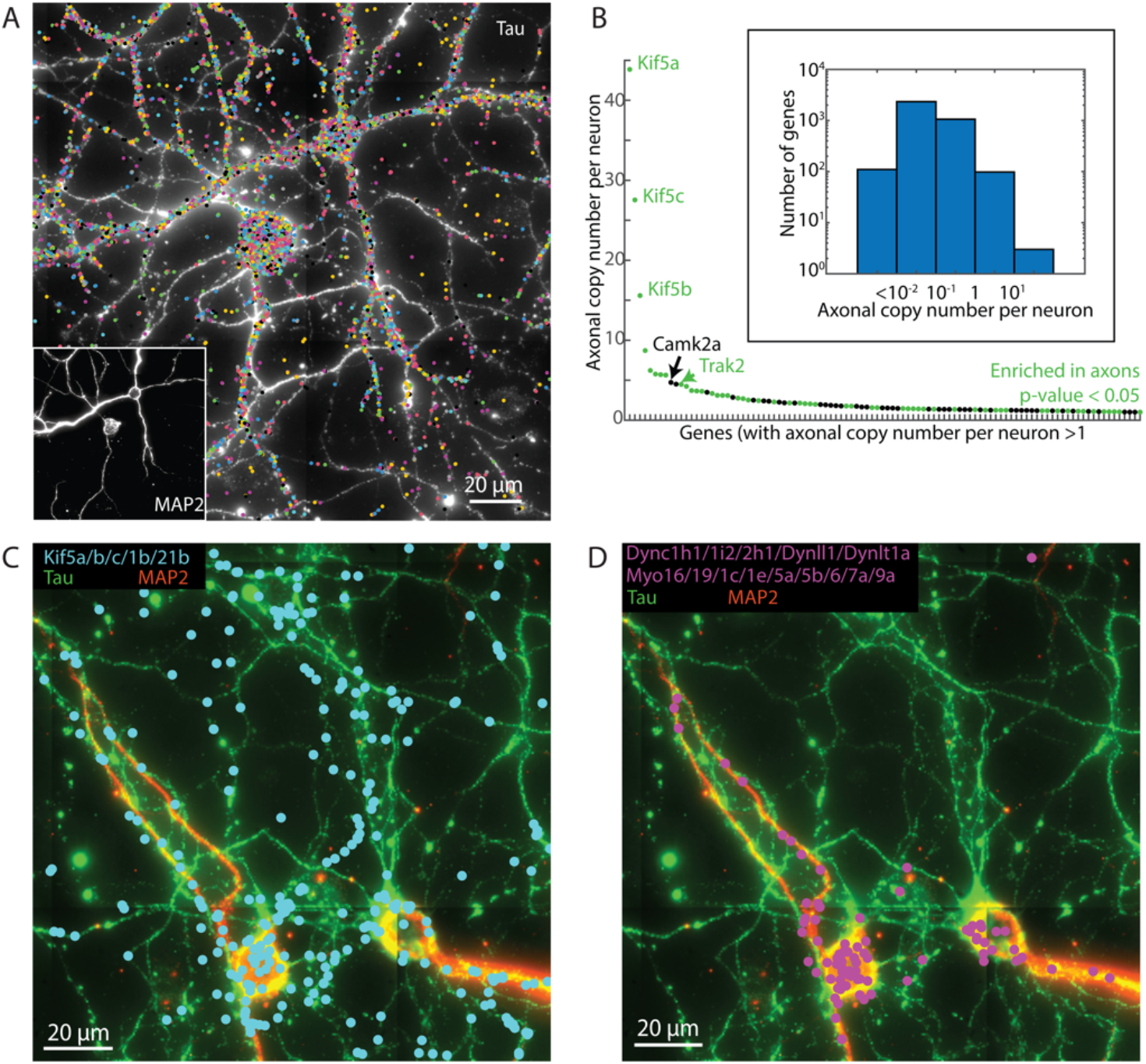
Identification and quantification of RNAs enriched in axons. (**A**) MERFISH image of 4209 genes in a sub-region of a 4 mm-by-4 mm imaged area. Each colored dot represents a single RNA molecule color-coded based on its gene identity. Tau fluorescence signal of the same area is visualized in the background as a black and white image. Scale bar: 20 µm before expansion. (inset) MAP2 fluorescence image of the same region. (**B**) Axonal copy number per cell of genes with more than 1 average copy of transcripts per neuron in axons (100 of the 4,209 genes passed this threshold), with genes rank-ordered based on their axonal mRNA copy number. Genes significantly enriched in axons are colored in green and the rest in black. (Inset) Distribution of axonal RNA abundances per neuron for all RNA species that were above the mean count of the blank control (3615 RNA species total). (**C**) Pseudo-colored image showing several kinesin mRNA species from a 4,209-gene measurement. Each RNA molecule is shown as a cyan dot. The Tau and MAP2 immunofluorescence signals are colored in green and red, respectively, in the background. (**D**) Pseudo-colored image of several dynein and myosin mRNA species from the same region as in (C). Each RNA molecule is shown as a magenta dot.

Notably, we observed exceptionally high abundance of some mRNAs encoding kinesins and kinesin-related proteins (*Kif5a, Kif5c, Kif5b, Trak2, Kif21b, Kif1b*) in axons, with a medium of 44, 28, 16, 4, 3, 2 copies per axon, respectively, (Fig. 2B, C). We observed similarly high axonal enrichments for these genes in the 950-gene measurements (Fig. S9). Interestingly, other motor protein genes, such as dynein- and myosin-related transcripts, were not enriched in axons, except for *Dynll1* (Fig. 2D).The Kif5-Trak-Miro complex is important for mitochondria transport and axonal maintenance (*25*) and has implications in hereditary spastic paraplegias (*26*) and amyotrophic lateral sclerosis (*27*). The extremely high abundance of *Kif5a-c* and *Trak2* mRNAs may provide a mechanism to locally produce kinesin motors for transport and maintenance of cargos, such as mitochondria and lysosomes, in axons.

The high spatial resolution of our measurements also allowed us to profile the spatial distribution patterns of RNAs within individual neurites. We next examined the sub-dendritic RNA distribution patterns in our 950-gene measurements. By considering the branching orders of different dendritic regions or separating dendrites into segments (each 25-µm long) of different distance to the soma, we observed gradual changes in the transcriptome profiles from proximal to distal dendrites (Fig. 3A). This profile change was similar between excitatory and inhibitory neurons, as reflected by the high correlation between individual genes’ mean distances along dendrites in excitatory neurons and those in inhibitory neurons (Fig. 3B). To further quantify the sub-dendritic RNA distribution patterns, we selected RNA species that passed a certain dendritic abundance threshold (>1 copy in dendrites per neuron) for analysis. We then determined, for each of these RNA species, the fraction of RNA molecules localizing to every 25-µm segments from proximal to distal dendritic ends, and normalized by z-score across genes for each 25-µm segment. We then performed hierarchical clustering analysis to group mRNAs based on their z-score profiles along the dendrites. We identified four major clusters of RNA species (red, orange, blue and purple groups in Fig. 3C), progressively more enriched in distal dendrites from the red group to the purple group. Similar results were observed in the 4,209-gene measurements (Fig. S10), The RNA groups most enriched in the distal regions of dendrites (blue and purple) encode many kinesin proteins, signaling molecules and structural components of the postsynaptic density (Fig. 3C and Fig. S10). Synapses are often localized to distal dendritic branches where locally synthesized proteins provides key support for synaptic functions (*10, 28, 29*). The mRNAs of synaptic proteins observed in distal dendritic regions could thus provide a means for local production of these proteins for the assembly and plasticity of synapses.

**Fig. 3.**
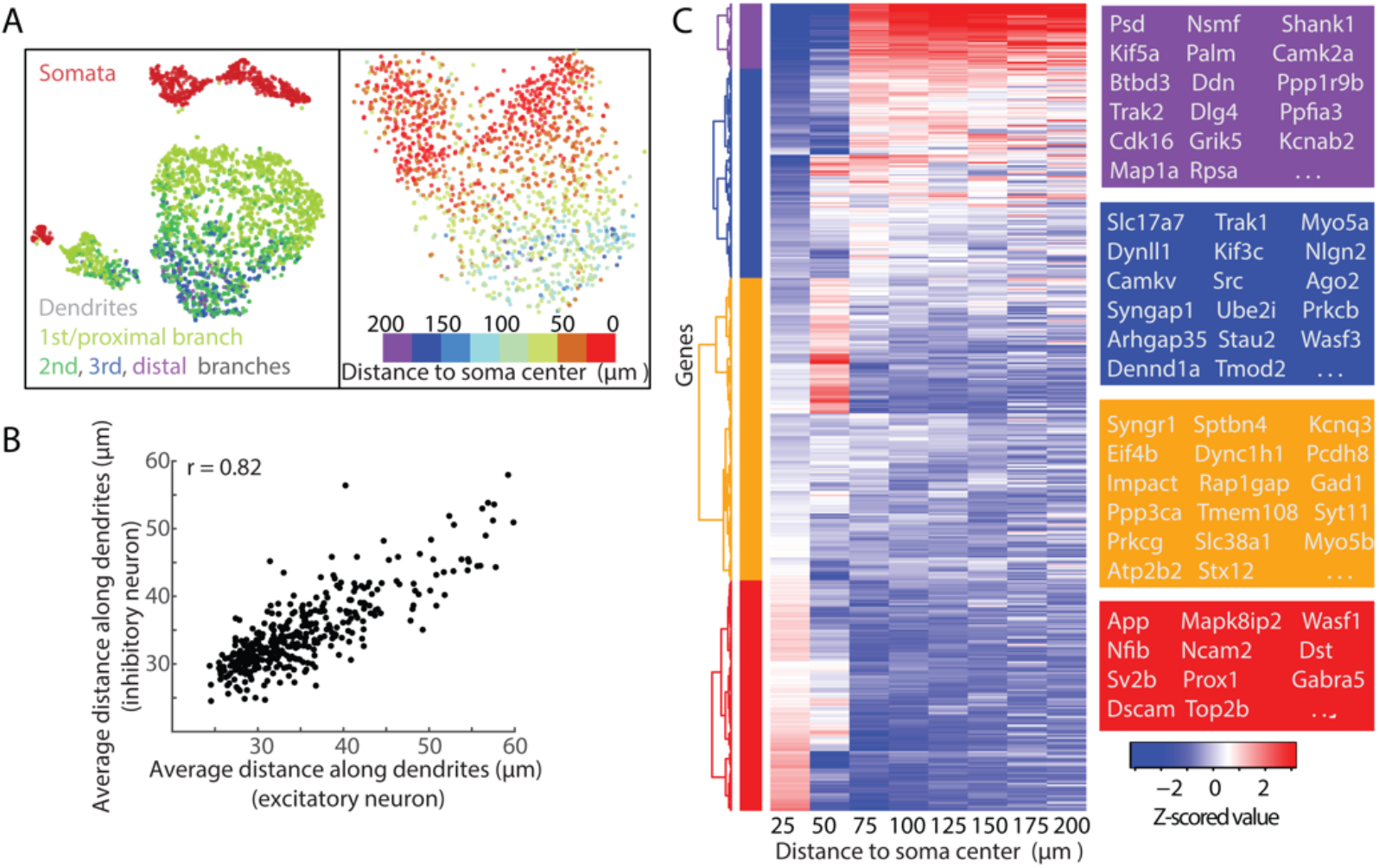
RNAs exhibit diverse distribution patterns along the proximal-to-distal axis of dendrites. (**A**) (left) t-SNE plot as in Fig. 1B but colored based on the subcellular compartment and the branching order of dendrites. (right) t-SNE plot for the transcriptome profiles of all dendritic segments (each 25-µm long) of excitatory neurons, with each segment colored based on its distance to the corresponding soma center. (**B**) Scatter plot of the average distance-to-soma along the dendrites for individual RNA species measured in excitatory neurons versus inhibitory neurons from the 950-gene measurements. Only RNA species with more than 1 dendritic copies per neuron in both excitatory neurons and inhibitory neurons are shown. The Pearson correlation coefficient (r) is 0.82. (**C**) Distribution of RNA molecules along dendrites for individual RNA species. Only RNA species with > 1 average copies in dendrites per neuron in the 950-gene measurements are shown here. Each row represents an RNA species. For each RNA species, we first computed the fraction of its RNAs localizing to each 25-µm segments along the dendrites, and then computed, within each 25-µm segment, the z scores of individual RNA species across all RNA species based on the aforementioned fraction quantification. The z-scores are shown in a heatmap according to the color scale shown on the right. Four major groups of RNA species were identified based on hierarchical clustering of the distribution patterns of individual RNA species. Boxes on the right showing examples genes in each of the four groups.

The copy numbers of most RNA species in the dendrites of individual cells are substantially lower than the number of synaptic sites per neuron, which poses a challenge for local translation at synapses, especially when translation from multiple RNA species are required in situations such as translation-dependent synaptic plasticity (*21, 30*). One possible way to overcome the paucity of RNAs is to cluster functionally related RNAs together spatially to create synergetic hotspots. To test whether some RNA species do have tendency to spatially cluster to form such hot spots, we examined spatial association or proximity between dendritic RNAs by accessing which RNA species were more likely to be nearest neighbors with each other (with a distance threshold of 1 µm, i.e. only nearest neighbors with distances <1 µm in 3D were considered). For this analysis, we chose to focus on the 950-gene measurements only, because the higher detection efficiency of these measurements provides a more accurate determination of spatial proximity between transcripts. We assessed the significance of spatial association or proximity by comparing this nearest-neighbor analysis with that of a randomization control where we permutated the RNA identities among detected RNA molecules while keeping the copy number of individual RNA species unchanged in individual dendritic branches (Fig. 4A).

**Fig. 4.**
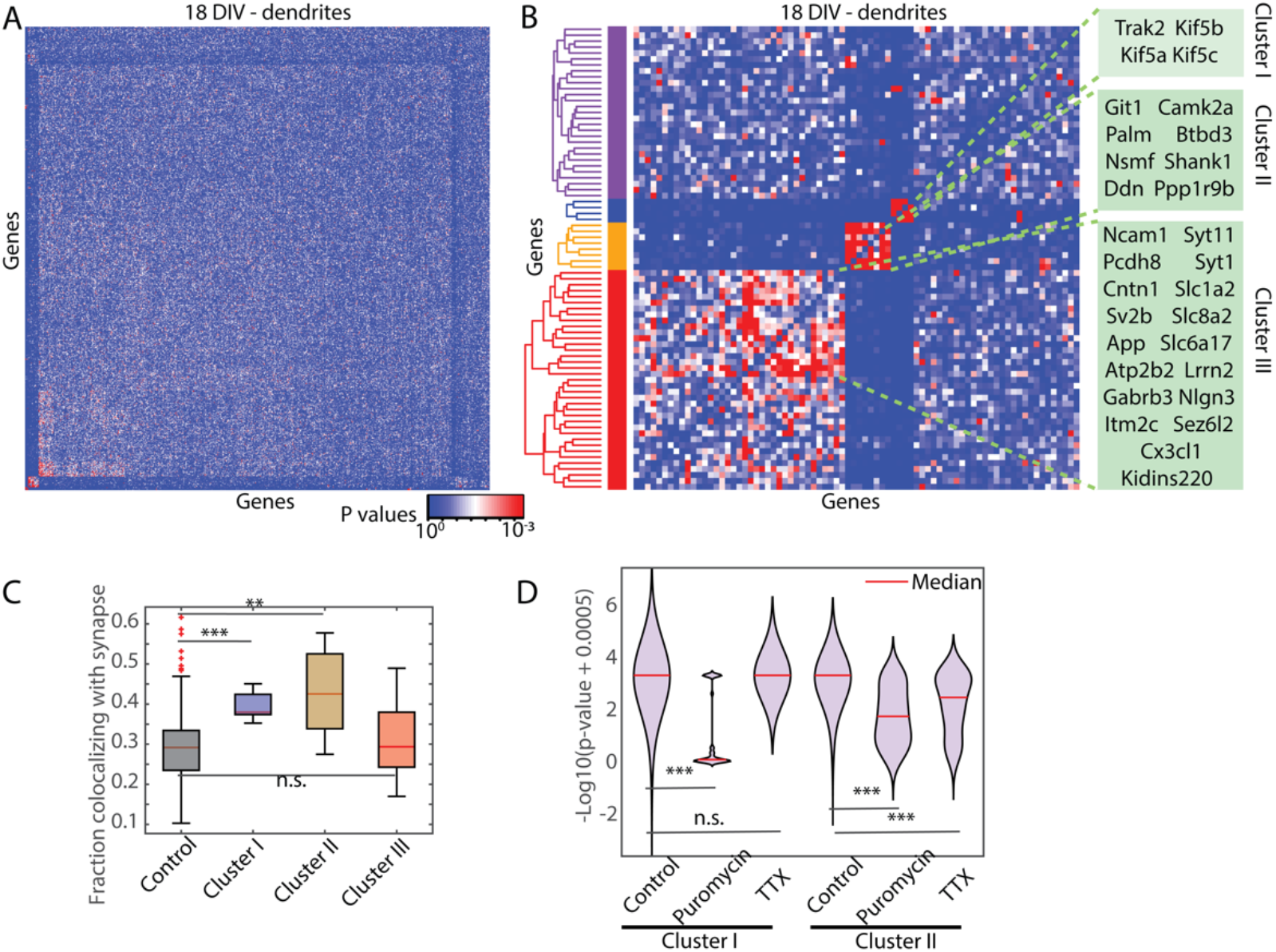
Distinct gene clusters exhibit spatial association of RNAs that are dependent on protein synthesis and differentially dependent on synaptic activity. (**A**) Heatmap of p-values of spatial association/proximity between RNA species in the dendrites of DIV 18 neurons. The p-value is calculated by comparing the frequency of an RNA species to be among the five nearest neighbors (and with a distance < 1 µm in 3D) of another species in the 950-gene MERFISH measurement with a permutation test while preserving the transcriptome contents in each dendritic branch. Each row represents the p-values of other RNA species to be enriched as nearest neighbors of a specific RNA species. (**B**) Same as (A) but for only those RNA species that had more than 3 other RNA species showing highly significant spatial association/proximity (p-value < 0.001). Hierarchical clustering separates these RNAs into multiple clusters. The boxes on the right list the RNA species three pronounced clusters. (**C**) Boxplot of the fraction of RNA molecules colocalized with synapses (with a distance threshold of 180 nm) for RNA species in each of the three clusters shown in (B), together with a control group that contains genes of similar dendritic abundance as the three clusters. The redline, box and whiskers of the boxplot represent median, the 25^th^ − 75^th^ percentiles, and the maximum range of data points not considered outliers (outliers shown as red crosses). (**D**) Violin plot of the −log10(p values) of spatial association/proximity between individual pairs of RNA species within clusters I and II (as defined in B) under different treatments. P-values are obtained by permutation test in neurites (2-4 biological replicates per condition). Tetrodotoxin (TTX) was added at 2 µM (>24 hrs); puromycin (Puro) was added at 100-200 µM (3 hrs). *** p-value < 0.001; ** p-value < 0.01; n.s.: p-value > 0.05 (two-sample two-sided Kolmogorov-Smirnov test).

Notably, we observed several clusters of genes with preferential spatial association or proximity among their transcripts in DIV 18 neurons – within each gene cluster, RNAs tended to have higher probability to be in spatial proximity with each other than with RNA species outside the cluster (Fig. 4A, B). When we examined RNA species with 3 or more significantly enriched nearest neighboring species, we observed three pronounced gene clusters (Fig. 4B). These clusters still persisted even when we performed a more conservative randomization that maintained RNA gradients for each RNA species along the proximal-distal dendritic axis (Fig. S11A), indicating that additional mechanisms beyond proximal-distal gradient along dendrites underlie these spatial association or proximity among RNAs. The spatial association patterns diminished in younger (DIV 10) neurons (Fig. S11B).

Next, we focused our analysis on these three clusters: a cluster represented by *Kif5a/b/c* RNAs that encode kinesin-1 subunits and *Trak2* RNA that encode trafficking kinesin protein 2 (Cluster I in Fig. 4B), a cluster represented by *Camk2a* RNA that encodes a calcium/calmodulin dependent protein kinase II subunit and *Shank1* that encodes a postsynaptic scaffolding protein (Cluster II in Fig. 4B), and a cluster represented by *App* RNA that encodes amyloid precursor protein and *Itm2c* RNA that encodes integral membrane protein 2C, a negative regulator of amyloid-beta peptide production (Cluster III in Fig. 4B). Cluster III contained many RNA species associated with outer mitochondrial membrane (6 out of 18 RNA species in this cluster and 11 out of 37 RNA species in its parent cluster highlighted by the red portion of the hierarchical tree in Fig. 4B are reported to be associated with outer mitochondrial membrane as measured by Apex-seq (*31*), whereas none of the RNA species in cluster II and only one of the RNA species in cluster I are reported to be mitochondria-associated).

To explore the mechanisms underlying the formation of clusters I and II, we first examined whether these RNAs are associated with synapses. To this end, we performed MERFISH imaging in combination with immunofluorescence to image these clusters of RNAs, as well as a control group of RNAs with similar dendritic abundance, together with MAP2 and pre-synaptic markers (a combination of VGLUT-1 and VGAT) for synaptic site identification. We identified synaptic sites as the overlapping regions of MAP2 positive dendrites and pre-synaptic signals in the immunofluorescence images. We found that, compared to the control RNA group, both clusters I and II were preferentially associated with synapses, whereas cluster III RNAs were not (Fig. 4C).

The fact that RNAs in clusters I and II formed two distinct clusters without detectable spatial association between the clusters (Fig. 4B) suggests that synaptic colocalization is unlikely to be the sole cause of the formation of these clusters. Interestingly, we found that inhibition of protein synthesis with puromycin significantly impaired the formation of both clusters I and II (Fig. 4D), indicating that the RNA association/proximity observed within these two clusters depended, at least partially, on protein synthesis and hence may have been facilitated by interactions between nascent peptides while these proteins were synthesized. For protein complexes made of distinct subunits, the interaction of nascent polypeptides during protein synthesis could bring the encoding mRNAs into proximity (*32, 33*). This mechanism could thus contribute to the formation of clusters I and II given our observation of their dependence on protein synthesis. Indeed, proteins encoded by *Kif5a/b/c* in cluster I are known to bind to each other to form the kinesin-1 protein complex. Protein products of several genes (*Camk2a, Shank1, Git1, Palm, Ppp1r9b*) in cluster II are synaptic proteins known to interact with the postsynaptic scaffolding protein PSD-95 (protein product of *Dlg4*) (STRING-db.org, (*34*)). These protein interactions could facilitate the clustering of their mRNAs. Consistent with this notion, even though not shown in Fig. 4B, we also observed significant spatial association/proximity of the mRNAs of *Camk2a, Palm* and *Ppp1r9b* with the *Dlg4* mRNA (p-value = 0.000, 0.002 and 0.05, respectively).

Notably, application of tetrodotoxin (TTX) to inhibit synaptic activity by blocking voltage-gated sodium channels impaired the formation of cluster II but not that of cluster I (Fig. 4D). Activity-dependent remodeling of post-synaptic density components can occur in a synergetic fashion, with specific groups of proteins accumulating or declining in synapses with similar kinetics (*35*). It is possible that RNAs in cluster II encode proteins that belong to one such module in response to synaptic activity. In addition, diverse RNA binding proteins accumulates at synapses upon synaptic activity stimulation (*36*). It is also possible that RNA localization by one or more RNA binding proteins (*3, 37, 38*) may contribute to the activity-dependent spatial proximity of RNAs in this cluster. Functionally, the spatial association of RNAs that we observed within these gene clusters could serve as synergistic platforms to facilitate local protein production and efficient assembly of proteins and protein complexes functioning in common pathways, and may offer cooperative translational control of specific multi-protein modules in response to synaptic activity.

In summary, we used MERFISH to quantify the copy numbers and spatial distributions of RNAs at genome scale inside individual neurons and sub-neuronal compartments with high spatial resolution. With quantitative, high-resolution spatial mapping of up to ∼4,200 mRNA species, our study created a subcellular atlas of the transcriptome in neurons, revealing quantitative abundances and enrichments for RNAs in axons and dendrites, as well as diverse sub-dendritic RNA distribution patterns. Furthermore, our measurements identified distinct clusters of RNA species with sub-micrometer-scale spatial association, the formation of which depended on protein synthesis and differentially on synaptic activity. These results provide a quantitative foundation for investigations of RNA localization and local RNA functions in neurons.

## Acknowledgement

We thank George Emanuel for advice and help with MERlin implementation for MERFISH decoding.

## Author contributions

G.W. and X.Z. designed the experiments. G.W. and J.R.M performed the experiments. G.W., C.-E.A., J.F. and A.W. performed data analysis. G.W., C.-E.A., J.F. and X.Z. wrote the manuscript with input from A.W. and J.R.M.

## Data and materials availability

All data and analysis software are available in the manuscript or available upon request.

## Supplementary Materials for

## Materials and Methods

### Neuronal culture

Briefly, hippocampus from embryonic day 18 (E18) mouse (timed pregnant C57BL/6J, the Jackson Laboratory, 000664 |Black 6) were isolated and digested with 0.25% trypsin–EDTA (1×) (T4549, Sigma) at 37 °C for 15 min, washed in HBSS solution (Life Technologies, 14175– 079) three times, and then transferred to the NB medium consisting of Neurobasal medium (Life Technologies, 12349–015) supplemented with 37.5 mM NaCl, 2% (vol/vol) B27 supplement (Life Technologies, 17504–044), 1% Glutamax (Life Technologies, 35050–061), and 1% penicillin–streptomycin (Life Technologies, 15140–122). After washing once with NB media, the tissues were pipetted up and down in NB medium until the tissues were mostly dissociated. Dissociated cells were then counted and plated onto poly-D-lysine–coated 18-mm coverslips (NeuVitro, GG-18–1.5-pdl) at 70,000 −100,000 cells per coverslip to achieve an optimal density for imaging and minimize bundling and overlap between cells. Cells were fixed at 18 or 10 days in vitro as indicated.

### MERFISH gene selection

#### 950-gene MERFISH measurement

RNA species include cell type marker genes (*Slc17a7, Gad1, Gad2, Prox1, Gfap, Pdgfra*), genes encoding motor proteins, and genes selected randomly from GO terms related to neurites and synapses.

#### 4209-gene MERFISH measurement

RNA species include cell type marker genes (*Slc17a7, Gad1, Gad2, Prox1, Gfap, Pdgfra*), genes previously observed to be present in axons (*22*–*24*) and genes selected from GO terms related to neurons.

### MERFISH experiment and decoding

#### MERFISH encoding probe design

MERFISH encoding probes for the 950-gene measurements were designed using a 32-bit Hamming-distance-4 (HD4), Hamming-weight-4 (HW4) code with 1,240 total barcodes. We randomly assigned the 1,240 possible barcodes to 950 RNA species and 290 blank barcodes. MERFISH encoding probes for the 4209-gene measurements were designed using a 49-bit HD4, HW4 code with 4,508 total barcodes. We randomly assigned the 4,508 possible barcodes to 4209 mRNAs and 299 blank barcodes. The 100-gene MERFISH libraries for detection efficiency comparison (Fig. S2) and the 110-gene MERFISH library for colocalization analysis with synaptic markers (Fig. 4C) were designed as described previously using 16-bit HD4, HW4 codes (*18, 20*). Each MERFISH encoding probe for each gene contains a 30-mer targeting sequences that is complementary to the target RNA and flanked by two readout sequences chosen randomly from the four possible readout sequences determined by the assigned hamming code of the target gene. The targeting sequences for a given RNA species were identified as previously described (*18, 20*), using mouse mm10 genome assembly and allowing up to 20-nt overlap between different targeting sequencing of an mRNA species, as described previously (*18*). Encoding probes were constructed as previously described (*18*). Dye labeled MERFISH readout probes complementary to the readout sequences on the encoding probes were synthesized and purified by Bio-synthesis, Inc.

#### Construction of oligo-conjugated antibodies

Donkey Anti-Mouse IgG, Donkey Anti-Rabbit IgG, Donkey Anti-Guinea Pig IgG and Donkey Anti-Chicken IgY secondary antibodies (Jackson Immuno, 703-005-155, 706-005-148, 711-005-152 and 715-005-150) were labeled with a copper-free click crosslinking agent using NHS-ester chemistry. Specifically, azide preservative was removed using a spin-column based dialysis membrane (Amicon, 100 kDa molecular weight cut off) according to the manufacturer’s instructions. DBCO-PEG5-NHS Ester (Kerafast) was diluted to a concentration of 5 mM in anhydrous dimethyl sulfoxide (DMSO) (Thermo Fisher Scientific). 2 µL of the solution was then combined with 100 µL of 1 mg/mL of the antibody in 1× phosphate-buffered saline (PBS). This reaction was incubated at room temperature for 1 hour and then terminated via a second round of purification using the Amicon columns as described above. Unique readout oligonucleotides with a 5’-acrydite, to allow cross-linking to the polymer gel, and a 3’-azide, to allow cross-linking to the DBCO-labeled antibodies, were ordered from IDT and suspended to 100 µM in 1× PBS. 20 µL of the appropriate oligonucleotide was then added to 100 µL of the DBCO-labeled antibodies (0.5 mg/mL). A third round of purification was performed before storing the product at 4°C. The average number of DBCO crosslinkers per antibody was determined via the relative absorption of the sample at 280 nm (antibody) and 309 nm (DBCO). On average the procedure described above produced ∼4 DBCO per antibody.

#### Neuronal sample preparation for MERFISH imaging

Neurons are fixed with 4% paraformaldehyde (PFA) (Electron Microscopy Sciences) in a cytoskeleton preserving solution (10mM MES, 138mM KCl, 3mM MgCl2, 2mM EGTA, 320mM Sucrose, pH6.1, Sigma) for 25 min, permeabilized with 0.25% (vol/vol) Triton X-100 (Sigma) in 1× PBS for 5-10 min, and washed with 30% (vol/vol) formaldehyde wash buffer in 2× SSC. MERFISH encoding probe staining, gel embedding, expansion and imaging were performed as described previously (*18*). Briefly, 15 µL hybridization buffer with encoding probes (∼2 nM per encoding probe for 950-gene library, 100-gene library and 110-gene library and ∼1 nM per encoding probe for 4,209-gene library; thanks to the background reduction and dilution by clearing and expansion, we observed better signals for higher probe concentration but the final concentration is limited by the throughput and cost for constructing these probes and solubility of probes in the hybridization buffer) and 3 µM of poly(dT) LNA anchor (IDT) was added to the surface of Parafilm (Bemis) and was covered with a cell-containing 18-mm coverslip. The hybridization buffer was composed of encoding wash buffer supplemented with 0.1% (wt/vol) yeast tRNA (Life Technologies), 1% (vol/vol) murine RNase inhibitor (New England Biolabs), and 10% (wt/vol) dextran sulfate (Sigma). Samples were incubated in a humid chamber inside a hybridization oven at 37 °C for 40 h. Cells then were washed with wash buffer twice (30 min each, 47 °C).

Immunostaining using oligo-conjugated antibodies can be performed either right after neuron fixation (before MERFISH) or after MERFISH encoding staining (before gel embedding). Briefly, cells were blocked at room temperature for 30 min in blocking buffer (4% (wt/vol) UltraPure BSA (Thermo Fisher Scientific) in 2× SSC supplemented with 3% (vol/vol) RNasin Ribonuclease inhibitor (Promega), 6% (vol/vol) murine RNAase inhibitor and 1 mg/ml yeast tRNA) and incubated with primary antibodies (Synaptic systems, 188 011, 314 006, 135 304, 106003, 131 005; Abcam, ab5392) in blocking buffer for 1 h at room temperature, and washed three times with 2× SSC for 10 min each. Samples were then incubated with oligo-labeled secondary antibodies in blocking buffer at a concentration of 4 µg/ml for 1 h at room temperature, then washed with 2× SSC three times for 10 min each. A brief fixation between MERFISH staining and immunofluorescence staining was introduced using 4% (vol/vol) PFA in 2× SSC for 5-10 min followed by three washes with 2× SSC. We observed slightly better immunofluorescence performance when it was carried out before MERFISH staining, and slightly better performance of MERFISH when immunostaining was carried out afterwards. Because of this, we applied MERFISH staining followed by immunofluorescence for all our MERFISH experiments of 950-gene measurements and 4209-gene measurements. For colocalization analysis with synaptic markers (Fig. 4C), we performed 110-gene MERFISH measurements, where we performed immunofluorescence prior to MERFISH encoding probe staining to better visualize synaptic signals, and we did not see obvious degradation of MERFISH signals for these measurements. In case of possible RNase contamination, murine RNase inhibitor (New England Biolabs) can be added to MERFISH and immunostaining buffers to preserve sample quality.

To de-crowd the RNAs, we embedded, expanded, and re-embedded the cultured cells using the reagents and protocols described previously (*18*), but with a modified low-salt buffer of 0.8× SSC instead of 0.5× SSC to tune down the expansion ratio to ∼2×.

Readout probe staining and MERFISH imaging was carried out as previously described with z-stack imaging of a 12-µm-depth volume with a step size of 1 µm (*18, 20*). We replaced the imaging buffer O_2_ scavenging enzyme with recombinant protocatechuate 3,4-dioxygenase (rPCO, OYC Americas) (*39*) to reduce possible RNase contamination. Our 40 mL of an imaging buffer comprised of 5 mM 3,4-dihydroxybenzoic acid (Sigma, P5630), 2 mM trolox (Sigma, 238813), 50 µM trolox quinone, 1:500 rPCO, 1:500 Murine RNase inhibitor, and 5 mM NaOH (to adjust pH to 7.0) in 2×SSC (*40, 41*).

#### MERFISH decoding

Images were aligned across multiple imaging rounds based on fiducial markers, high-pass filtered and deconvolved, and normalized for intensity variation across multiple rounds. 100-gene and 110-gene MERFISH images were decoded as previously described (*18, 20*). Images for 950-gene and 4209-gene MERFISH images were decoded using MERlin package (*19*) (GitHub/ZhuangLab/MERlin), which is optimized for MERFISH libraries with high complexity. Briefly, images were decoded in 3-D using hamming distance threshold of 0.77 (the maximum distance for barcodes with 2-bit errors), where adjacent voxels with the same barcodes were considered the same molecule. Afterwards, only barcodes with 2 voxels or more were kept and adaptive thresholding strategies were adopted to construct the final list of detected transcripts with 5% misidentification for the 950-gene MERFISH and 10% misidentification rate for the 4209-gene MERFISH library (<0.2 average errored copy per RNA species per cell). We note that generally, a lower mis-identification rate is accompanied by a lower detection efficiency of transcripts and hence, we chose a higher misidentification rate in 4209-gene measurements to partially compensate for the reduction in detection efficiency at high RNA density.

### Annotation of dendritic and somatic regions

Somata and dendrites were segmented, and the dendrites were annotated manually using a custom-written MATLAB script. The annotation of dendritic and somatic regions was performed by experienced individuals based on several cell marker FISH signals (*Slc17a7, Gad1*/*2, Pdfgra, Gfap*) and the MAP2 and Tau immunofluorescence signals. The following general rules were used: 1) Somata were identified by their high density of mRNAs, detectable immunofluorescence signals of both MAP2 and Tau, and recognizable neuronal cell body shape. When multiple somata were located adjacent to each other, boundaries were drawn at where the mRNA density drops suddenly. 2) We assigned dendrites to their corresponding somata by tracing their MAP2 signals back to the originating somata. When two dendrites overlap, we removed the region of intersection from the segmentation unless one has overwhelmingly higher density than another, in which case, we assigned the region to the higher-density dendrite. We traced and annotated every dendritic branch until they can no longer be confidently matched to a given cell. 3) If a dendrite appeared likely to extend from more than one soma, we annotated it only when there was a clear decreasing gradient of mRNA density from only one soma or if the cell type markers (for excitatory/inhibitory neurons) in the dendrites match only one of the somata. 4) For cells with weak MAP2 and Tau immunofluorescence signals, we annotated somata based on the density of mRNAs and annotated their protrusions based on the gradient of mRNAs. 5) For cells whose neurites overlapped significantly with other cells or cells with markers for glia and oligodendrocytes, we did not annotate neurites but only recorded the soma boundaries. These cells were not included in analysis regarding dendritic transcriptome properties.

### t-distributed stochastic neighbor embedding (t-SNE) analysis

After dendrites and somata were manually annotated, we kept only neuronal cells with strong *Gad2* or *Slc17a7* expression (>50 copies per cell for 950-gene measurement or >10 copies per cell for 4,209-gene measurement) and with low *Gfap* and *Pdgfra* mRNA expression (<10 copies per cell for 950-gene measurement or <5 copies per cell for 4,209-gene measurement). t-SNE analysis was performed on all annotated somata and dendritic branches based on RNA abundance measured in the 950-gene measurements. Every annotated dendritic branch was used in the analysis, and as a result there were many more dendritic points than somatic points in the t-SNE plots.

Since we did not observe strong batch effects between different biological replicates, we did not perform batch correction. For t-SNE visualization in Fig. 1B and Fig. 3A (left), we first performed CPM (count per million) normalization on the MERFISH counts per dendritic or somatic compartment. We then normalized variance as a function of expression magnitude to identify 382 overdispersed genes (generalized additive model fit with k = 5, alpha significance threshold = 0.2). We perform dimensionality reduction using principal component analysis (PCA), restricted to the 5 principal components (PCs) with the highest eigenvalues, and finally visualize using a 2-D t-SNE embedding (perplexity=50) (*42*). For t-SNE visualization in Fig. 3A (right), similar procedures were performed on the 25-µm dendritic segments from proximal to distal ends of excitatory neurons. We got 47 overdispersed RNA species and used 10 PCs instead for analysis.

### Calculation of dendrite-to-soma ratio

From the neuronal cells (see details above), we next removed neurons that did have enough dendritic areas annotated by only keeping cells with dendritic area that were no less than 50% of its somatic area. The apparent dendrite-to-soma ratio was calculated for each gene in each dataset by dividing the RNA counts in traced and annotated dendritic regions by the RNA counts measured in somatic regions. For 950-gene MERFISH measurements, since the detection efficiency in soma was roughly 45% of that in dendrites, and on average ∼50% of dendritic barcodes were included in the annotated dendrites, these two factors approximately balanced out and the true dendrite-to-soma ratio should approximately equal to the apparent dendrite-to-soma ratio. For 4,209-gene MERFISH measurements, we further calibrated the dendrite-to-soma ratio based on the detection efficiency difference between 4,209-gene MERFISH measurements and 950-gene measurements.

### Differential expression analysis during neuron maturation

Differential gene expression was done by using the Bioconductor package DESeq2 (*43*) on MERFISH counts in the dendritic or somatic compartments of excitatory neurons (marked by Slc17a7). P-values were attained by the Wald test and corrected for multiple testing using the Benjamini and Hochberg method.

### Identification of RNAs enriched in dendrite

To understand the baseline of dendrite-to-soma ratios, we utilized the 4,209-gene scale measurement, and selected somatic genes that are annotated in GO:0043025 neuronal cell body, but absent from GO terms related to neurites and synapses. This somatic gene list contains 268 genes, and the median dendrite-to-soma ratio is 0.16.

To identify RNA species enriched in dendrites, we performed a randomization test where the RNA identity labels were randomly permutated 5,000 times within each neuron while keeping the localizations of individual RNA molecules and the total copy number of each gene unchanged. The distribution of dendritic abundance of an RNA species was computed based on the permutation results. An RNA species was classified as being enriched in dendrites when the measured dendritic count was above the 95% quantile (i.e. p-value < 0.05) of the distribution determined by the above random permutations, and the fold of enrichment was determined as the ratio of the measured count divided by the average count in permutated results.

### Identification of RNAs present in axons

Axonal RNAs were classified based on MAP2 and Tau immunofluorescence images. First images were aligned using affine transformation based on the intensity profiles and binary masks were generated for MAP2 and Tau images. Then for each barcode, its minimum distances to MAP2- and Tau-positive regions were computed. Axonal RNAs were defined as RNAs that were closer to Tau-positive regions than to MAP2-position regions, that were within threshold distance (5 µm) to Tau-positive regions, and that had higher Tau signals than MAP2 signals at their locations. These criteria helped identify RNAs localizing to the distal arbors of axons which could have dim Tau signals.

### Identification of RNAs enriched in axons

To identify RNA species enriched in axons, we performed randomization tests where the RNA identity labels were randomly permutated 5,000 times within each field of view while keeping the localizations of individual RNA molecules and the total copy number of each gene unchanged. The distribution of axonal abundance of an RNA species was computed based on the permutation results. An RNA species was classified as being enriched in axons when the measured axonal count was above the 95% quantile (i.e. p-value < 0.05) of the distribution determined by the above random permutations, and the fold of enrichment was determined as the ratio of the measured count in axons divided by the average count in permutated results.

### Calculation of gradients along the proximal-distal axis of dendrites

Neurons with dendritic area that were no less than 50% of its somatic area (as defined above) were included for gradient analysis. Dendrites were traced using a custom-written MATLAB script. To determine the distance of each RNA molecule to the corresponding soma along the dendrite, we first computed the path along which the dendrite extends. To do this, we converted the dense RNA distribution in dendrites to a graph representation with each node corresponding to a 2.5 µm by 2.5 µm region occupied by RNAs. The adjacency matrix denoting undirected edges between nodes were assigned to the nearest neighboring node of each node or between nodes less than 5 µm away in the physical space. In distal dendritic regions where nodes were very sparse (more than 5 µm away from each other), we iterated the nearest neighbor assignment until the connectivity allowed us to trace every distal node back to the most proximal node. The shortest path between the end nodes (i.e. the distal end of a dendritic branch and the proximal dendritic branching point) was computed using the MATLAB function shortestpath and was used as the path along which the dendrite extends. The distance of an RNA to the soma center was calculated along this path.

The hierarchical clustering was performed based on the distribution of RNAs located within 150 µm from the soma center. First we calculated the fraction of RNAs localizing to every 25-µm segment along the dendrite; then we normalized the value in each segment by z-score across different RNA species in the same segment, based on which we performed hierarchical clustering with Euclidean distance and Ward linkage method. Four major groups were identified this way and within each group, and the clustering tree was reordered based on the average distance to soma center for a given RNA species, using mean as the function for weights agglomeration and maintaining the constraints on the tree.

### Gene ontology (GO) analysis

GO term enrichment was determined using PANTHER (http://pantherdb.org/, (*44, 45*)). For GO term analysis of the dendrite-enriched RNAs, enriched GO terms were determined for the RNA species that were consistently and substantially enriched in dendrites in all datasets (fold of enrichment > 1.5 and p-value < 0.05). We further compared the list to enriched GO terms identified for the rest of the RNAs in our measurements, and only the GO terms uniquely present in the former group were kept.

### Nearest neighbor analysis and enrichment test

We first identified the nearest neighbors of each RNA (termed as RNA-x here) within a small distance threshold (1 µm), excluding RNAs of the same species as RNA-x. We then recorded the top 5 most adjacent RNAs for a given RNA and combined the statistics for all RNAs of the same species to get the neighbor count profiles of this RNA species. To get the enrichment p-values, we randomly permutated the gene labels in a measurement, while preserving the spatial localizations of individual RNA molecules. For example, for permutation within a single dendrite, all RNA labels within the same dendritic branch were permutated for 2000 times and the neighbor profiles reanalyzed. The p values were calculated by comparing the measured result of RNA-RNA neighbor pair count to the neighbor count distributions in the permutation test. For permutation test for neurites (in regions outside somata), the permutation was carried out among all RNAs localized in neurites within the same optical field-of-view (∼200 µm by 200 µm). For permutation test preserving dendritic gradients, dendrites were first segmented into a 5-µm-long segment and the permutation was carried out among all RNAs localizing within the same segment.

The heatmaps in Fig. 4 and Fig. S11 were constructed based on the p-values obtained from the permutation test. The RNA species in the row and column were reordered according to the clustering tree from hierarchical clustering of genes in each row with Euclidean distance and Ward linkage method.

### Visualization of the MERFISH data

We created a web application to enable visualization and interactive exploration of the MERFISH data. This allows selected genes of interested to be highlighted using different colors and the spatial distribution patterns to be visually explored. The segmentation of cells and cell processes can also be visualized as convex hulls simultaneously with the selected RNAs. RNAs can be further overlaid on a composite MAP2/Tau staining image with MAP2 and Tau in different colors. The opacity of the staining image and size of mRNA spots can be adjusted to facilitate visual exploration.

This interactive web-application was implemented using the open-source data explorer, which we termed MERmaid. MERmaid uses the deck.gl WebGL-powered framework and React to render large-scale datasets such as millions of RNA molecules with high computational performance. MERmaid allows complex visualizations such as highlighting multiple genes overlaid onto background staining images by using multiple visualization layers. MERmaid supports input of gzipped csv format data with each row as a single gene with x, y coordinates along with other metadata information related to cell segmentation if available. The source code for the MERmaid data explorer is available through Github at https://github.com/JEFworks/MERmaid.

## Supplementary Figures

**Fig. S1.**
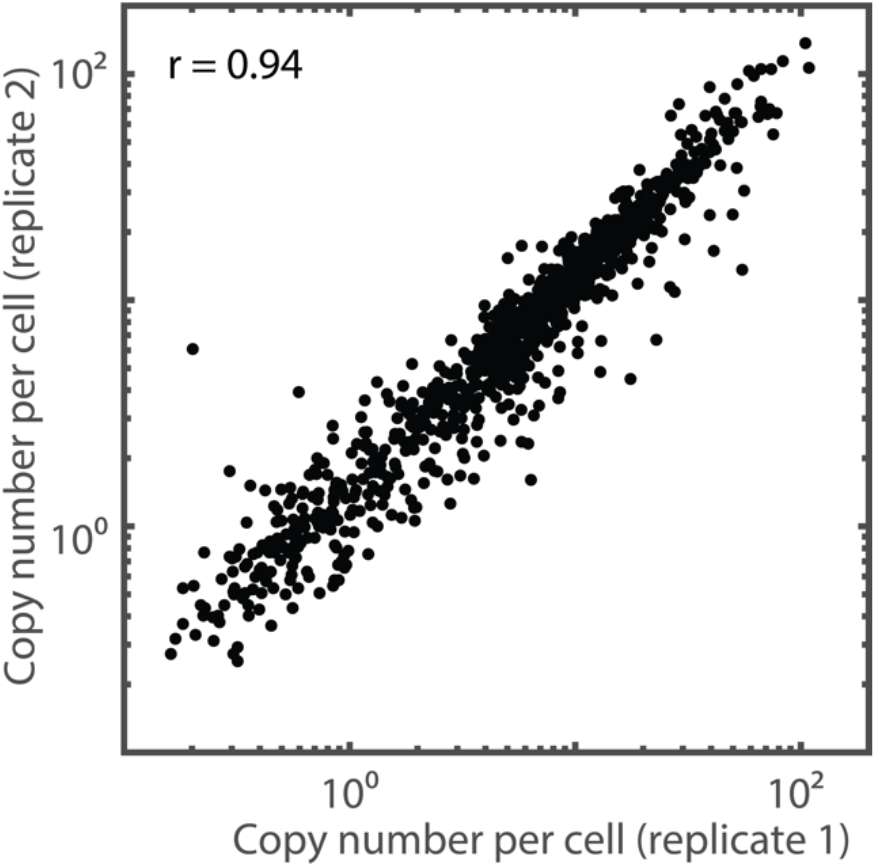
Comparison between replicates of 950-gene MERISH measurements. The scattered plot shows the average RNA copy numbers per cell of individual genes detected in one biological replicate versus those detected in another replicate. The Pearson correlation coefficient (r) is 0.94.

**Fig. S2.**
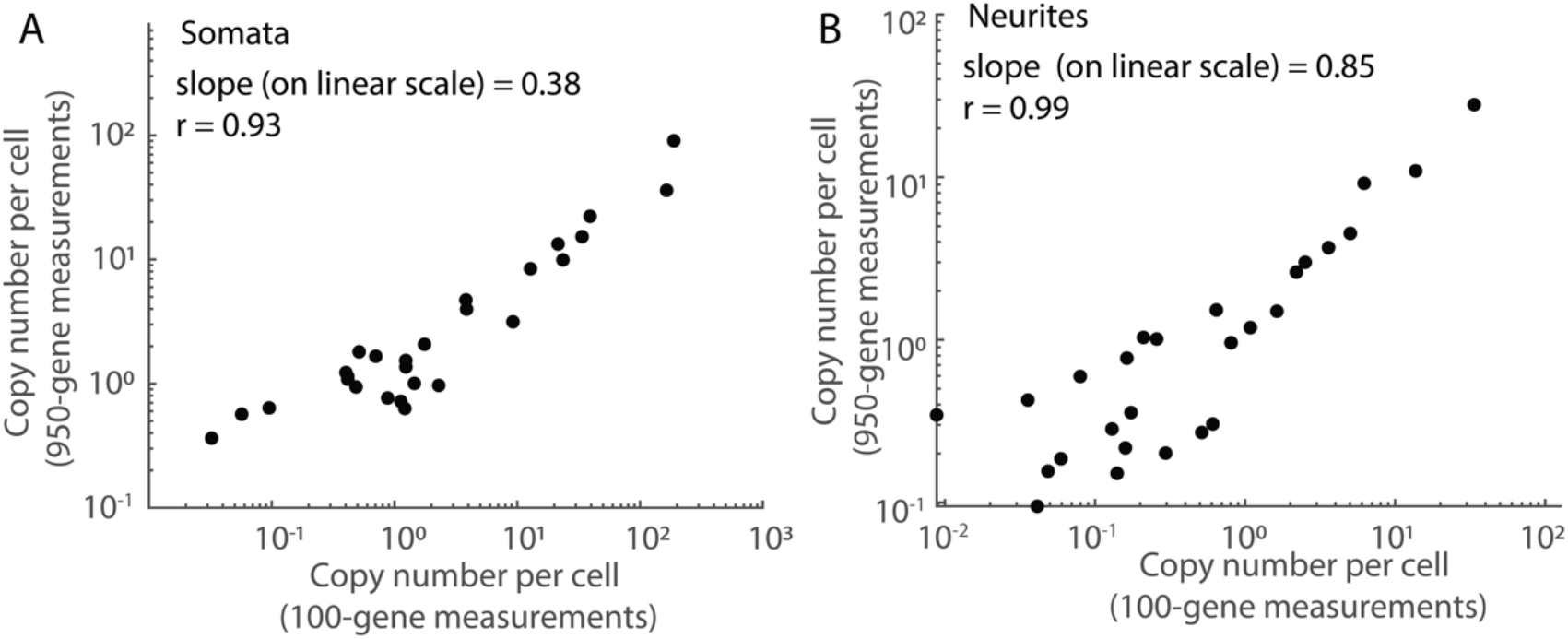
Comparison between 950-gene and 100-gene MERFISH measurements. (**A**) The average RNA copy numbers per soma determined in 950-gene MERFISH library (4 replicates) versus those determined in ∼100-gene MERFISH measurements (2 replicates). The Pearson correlation coefficient (r) is 0.93 and the linear regression slope of the data is 0.38 (both determined with the data plotted on the linear scale). Only RNA species with copy number per cell greater than 1 was considered in the linear regression. (**B**) Same as (A) but for RNAs detected in neurites (r = 0.99, slope = 0.85, determined as in (A)).

**Fig. S3.**
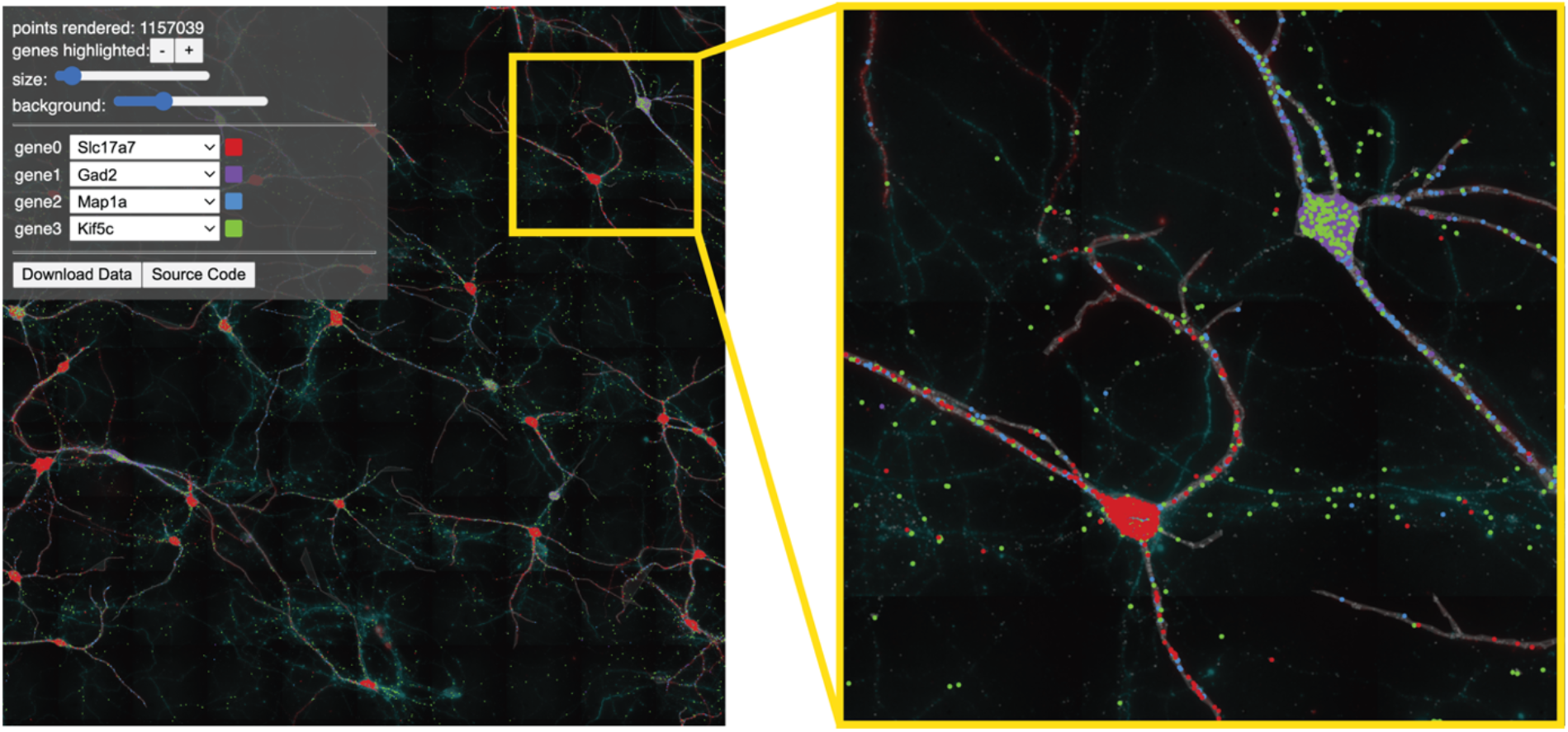
MERmaid data explorer enables interactive visualization and exploration of MERFISH data. (left) Screenshot of the data explorer. A menu allows users to select multiple genes of interest simultaneously and display them in different user-defined colors. The size of individual mRNA spots and the opacity of the background staining image may also be adjusted by the user. (right) A zoomed in region. We note that the RNA spots with the same x, y values are overlaid according to their z values with high-z spots masking low-z spots (as a result some of the blue and green RNA spots are masked). Adjusting opacity (set at 100% in the figure) will allow spots with lower z-values to show up.

**Fig. S4.**
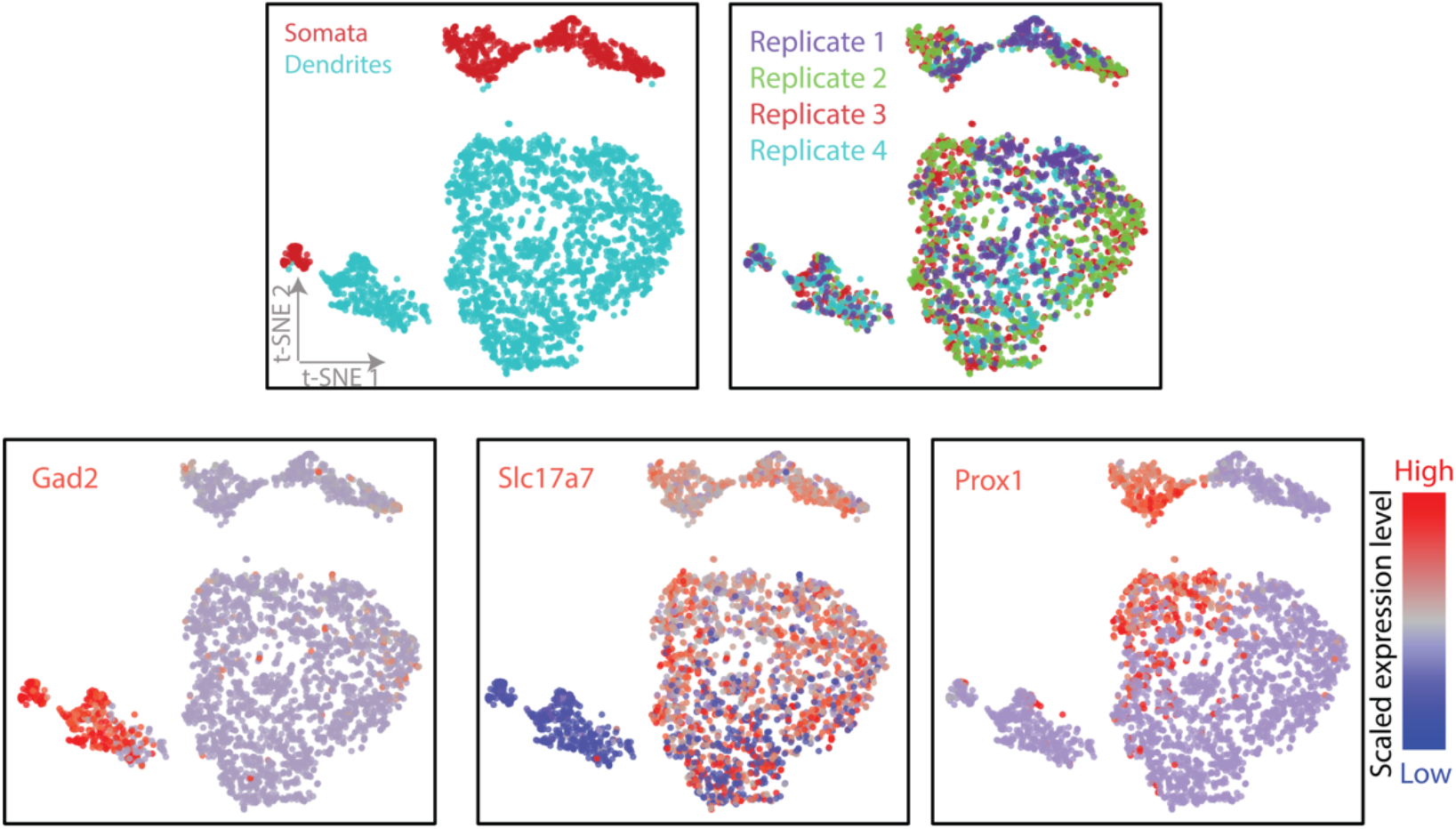
t-SNE plots for all individual somatic and dendritic compartments based on their transcriptome profiles. (Top row) Left: Replotting Fig. 1B for reference. Right: Different replicates show similar distributions on the t-SNE plot. Each spot is colored by which biological replicate it comes from. (Bottom row) Expression of some cell type markers. Left: Each point in the t-SNE is colored by *Gad2* expression level. Middle: Each point in the t-SNE is colored by *Slc17a7* expression level. Right: We also observed other cell type markers subdividing major cell types (e.g. *Prox1* in excitatory neurons). The expression level of the marker genes is indicated in colors according to the color scale shown on the right.

**Fig. S5.**
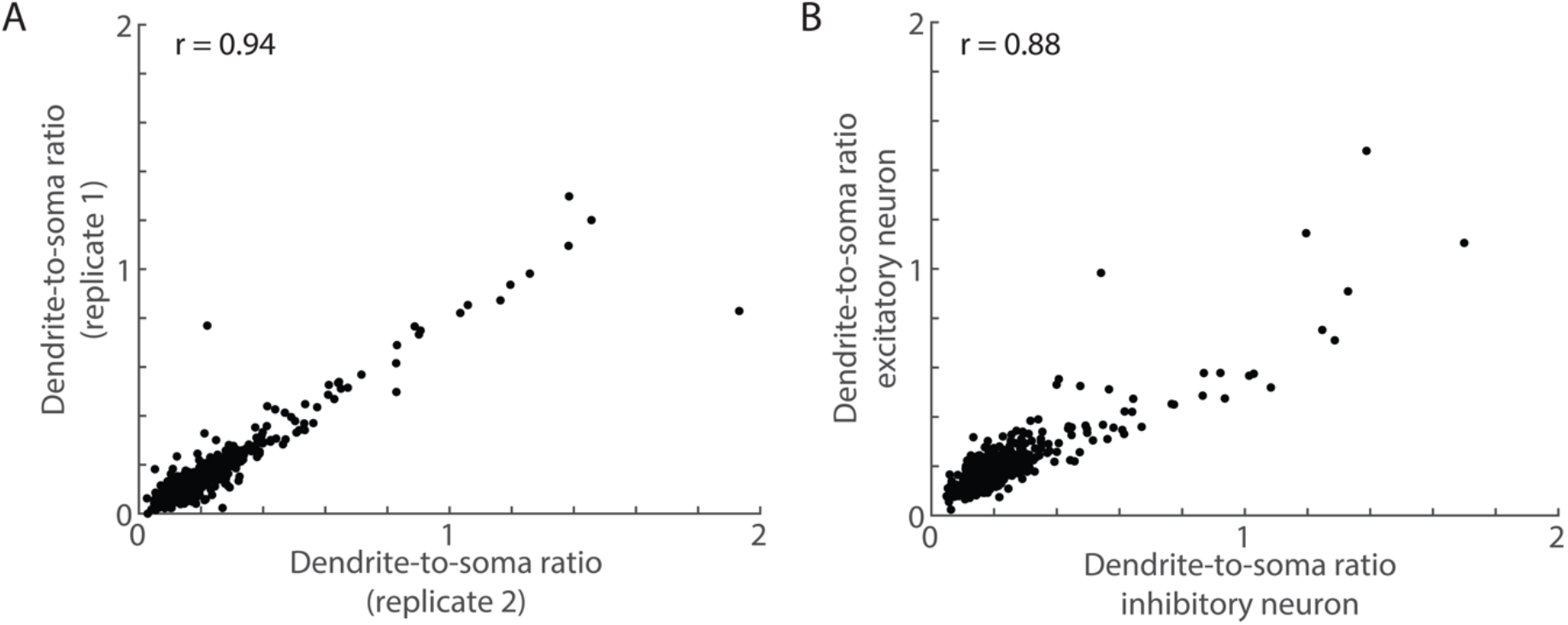
Comparison of dendrite-to-soma ratios between replicates and between excitatory and inhibitory neurons. (**A**) Scatter plot of dendrite-to-soma ratios of individual genes from measurements conducted on two biological replicates of the 950-gene measurements. Each dot represents an RNA species. The Pearson correlation coefficient (r) is 0.94. (**B**) Scatter plot of average dendrite-to-soma ratios measured in excitatory neurons versus inhibitory neurons from all four biological replicates. Only RNA species with more than 0.1 copies per neuron in dendrites in both excitatory neurons and inhibitory neurons are included. The Pearson correlation coefficient (r) is 0.88.

**Fig. S6.**
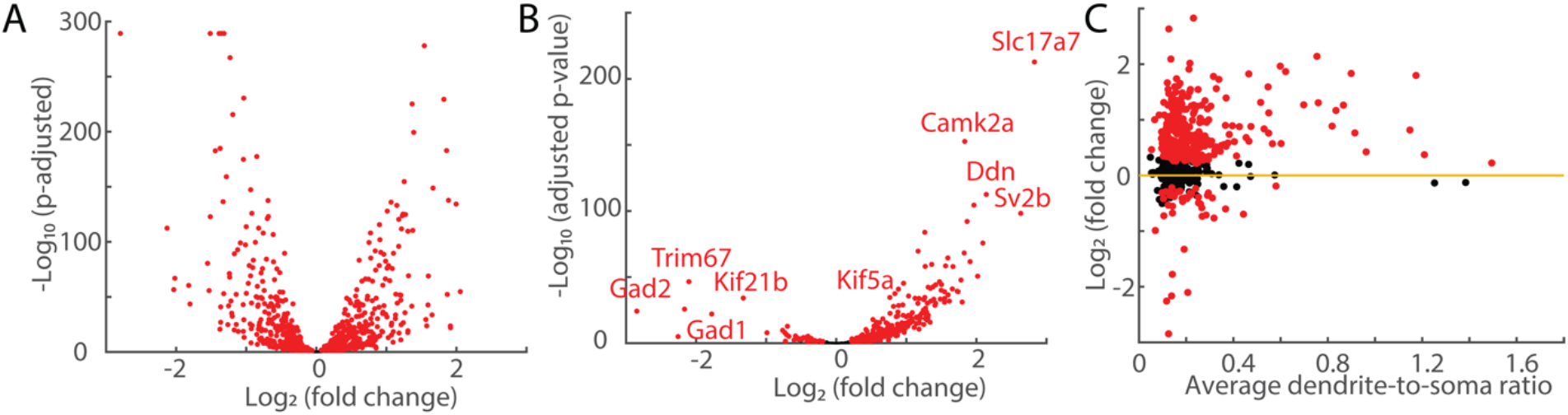
Changes in RNA expression in somata and dendrites upon neuronal maturation. (**A, B**) Quantification of changes in enrichment of RNA species upon neuronal maturation using DESEQ2. Volcano plot showing changes in enrichment of RNAs in the somata (A) and dendrites (B) between DIV18 and DIV10 excitatory neurons. The x-axis displays the fold change between DIV18 and DIV10 for individual RNA species and the y-axis displays the corresponding adjusted p values. RNA species with adjusted p values less than 0.05 are labeled in red and others in black. Some of the significantly enriched/de-enriched RNA species are labeled at the corresponding locations to serve as examples. (**C**) Scattered plot of Log2 fold change of dendritic RNA counts between DIV18 and DIV10 excitatory neurons versus dendrite-to-soma ratio in DIV 18 excitatory neurons for individual RNA species. RNA species with adjusted p values (DESEQ2) less than 0.05 are labeled in red and others in black.

**Fig. S7.**
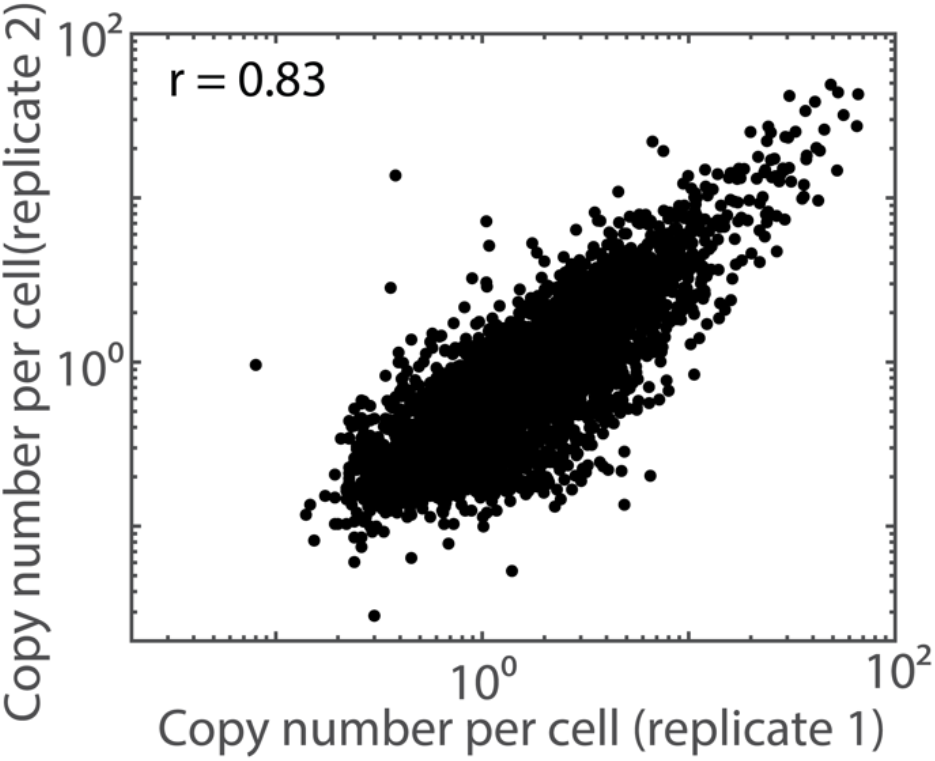
Comparison between replicates of 4209-gene MERISH measurements. The scattered plot shows the average RNA copy numbers per cell of individual genes detected in one biological replicate versus those detected in another replicate. The Pearson correlation coefficient (r) is 0.83. Only transcripts in annotated neurons were included in the analysis.

**Fig. S8.**
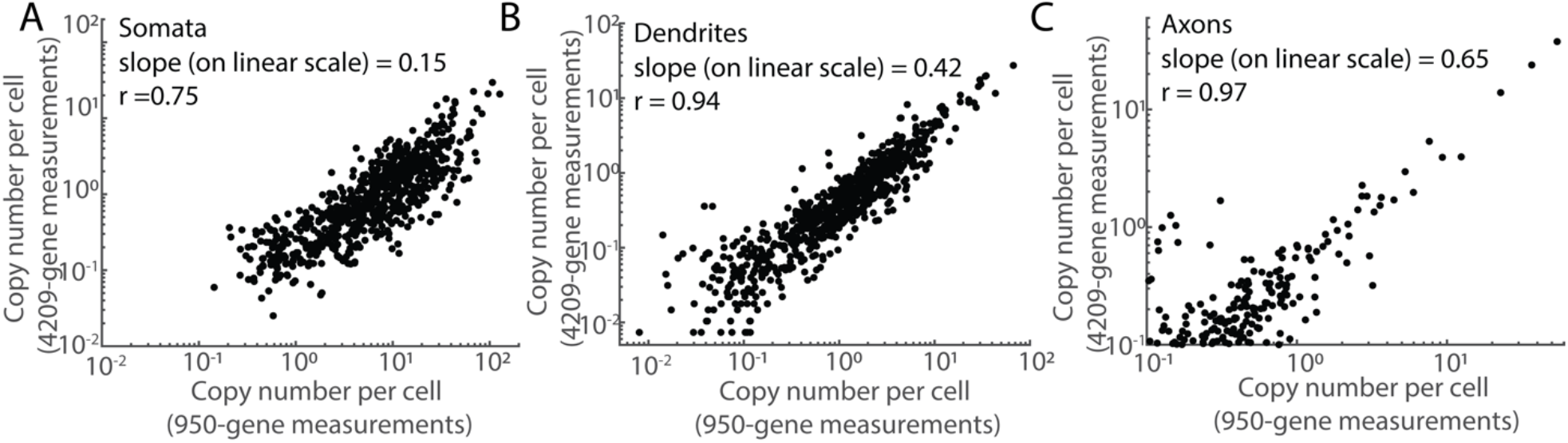
Comparison between 4209-gene MERFISH measurement and 950-gene MERFISH measurements. **(A)** The somatic RNA copy numbers per cell determined in 4209-gene MERFISH measurements (2 replicates) versus those in 950-gene MERFISH measurements (4 replicates). The Pearson correlation coefficients (r) is 0.75 and the linear regression slope of the data is 0.15 (both determined with data plotted on the linear scale). Only RNA species with copy number per cell greater than 1 is considered for the slope determination. **(B)** The dendritic RNA copy numbers per cell determined in 4209-gene MERFISH library versus those in 950-gene MERFISH measurements. The Pearson correlation coefficient and linear regression slope (r = 0.94, slope = 0.42) were determined as in (A). Only RNA species with dendritic copy number per cell greater than 0.5 is considered for the slope determination. **(C)** The axonal RNA copy numbers per cell determined in 4209-gene MERFISH library versus those in 950-gene MERFISH measurements. The Pearson correlation coefficient and linear regression slope (r = 0.97, slope = 0.65) were determined as in (A). Only RNA species with axonal copy number per cell greater than 0.5 is considered for the slope determination.

**Fig. S9.**
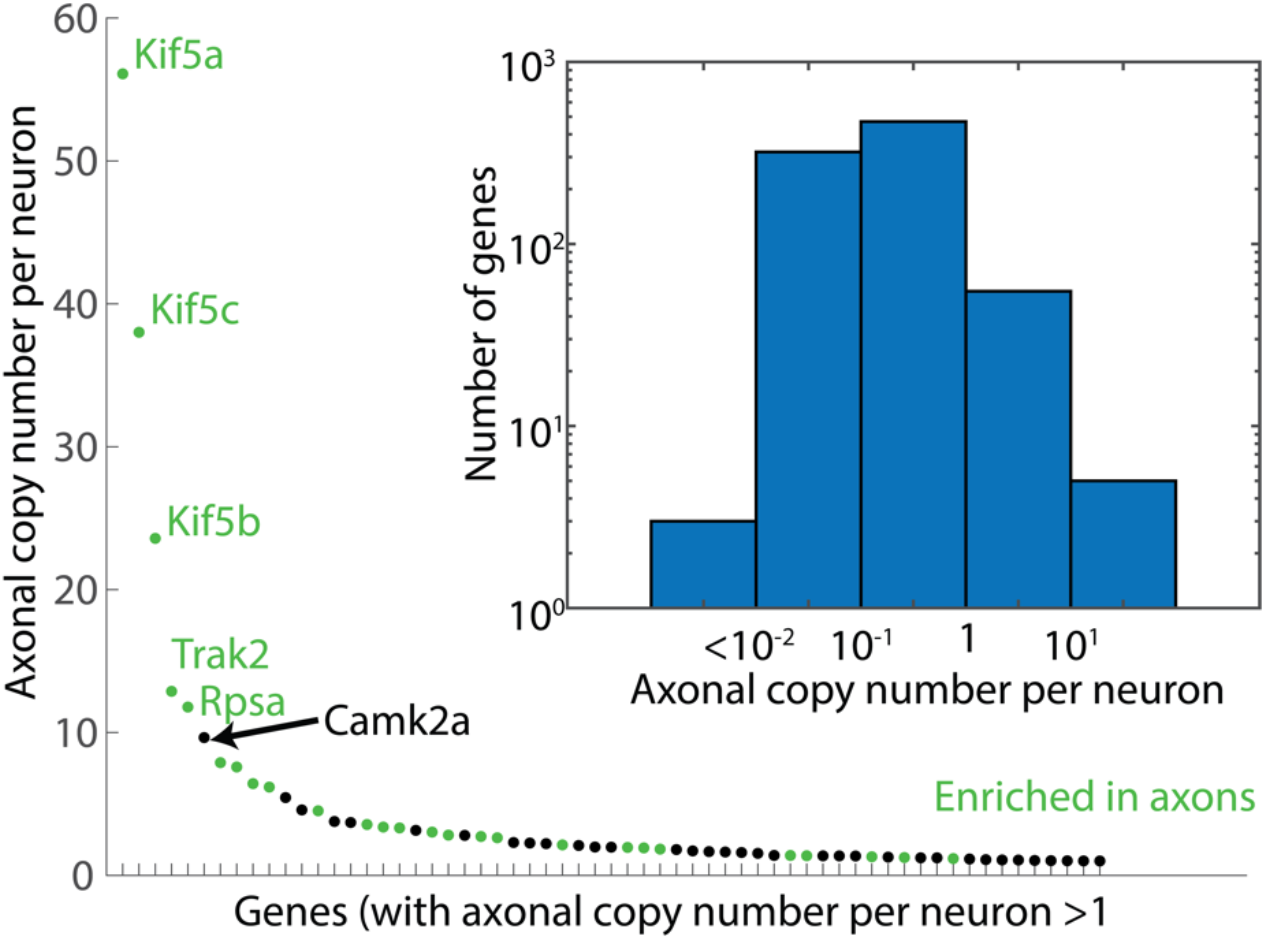
Axonal RNA analysis for the 950-gene MERFISH measurements. Axonal copy number per cell of genes with more than 1 average mRNA copy per neuron in axons, with genes rank-ordered based on their axonal mRNA copy number. 61 of the 950 genes have mRNA copy number greater than 1. Genes significantly enriched in axons (as compared to somata and dendrites) are colored in green and the rest in black. (Inset) distribution of axonal RNA abundances per neuron for all RNA species above the mean count of blank control (853 RNA species).

**Fig. S10.**
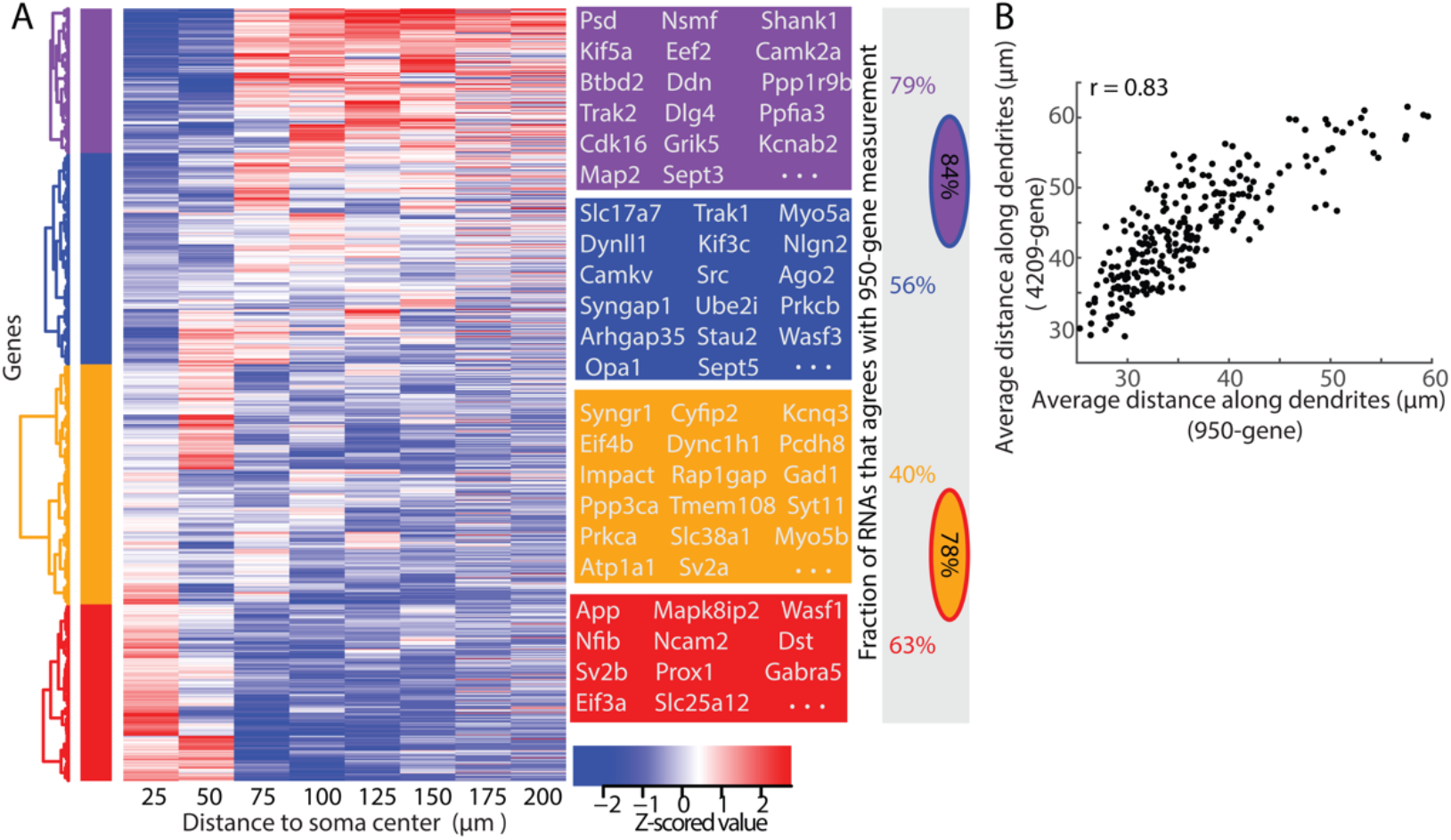
RNA distributions along the proximal-to-distal axis of dendrites measured in 4,209-gene MERFISH measurements. (**A**) Left: Distribution of RNA molecules along dendrites for individual RNA species, as described in Fig. 3C, but for the 4,209-gene measurements. Middle: Boxes showing examples of RNA species in each of the four major groups identified by hierarchical clustering. Right: the fraction of RNA species in each group whose cluster identities agree with the identities determined in 950-gene MERFISH measurements. Only RNA species included in both 950-gene and 4209-gene libraries were considered in determining these fractions. The oval circle (with the boundary and the shading in two different colors) shows the fraction quantification when the two groups of the corresponding colors are combined. (**B**) Scatter plot of the average distance-to-soma along the dendrites for individual RNA species measured in 4,209-gene MERFISH measurement versus 950-gene measurements. Only RNA species with more than 1 dendritic copies per neuron in 950-gene measurement and 0.6 dendritic copies per neuron in 4,209-gene measurement are included.

**Fig. S11.**
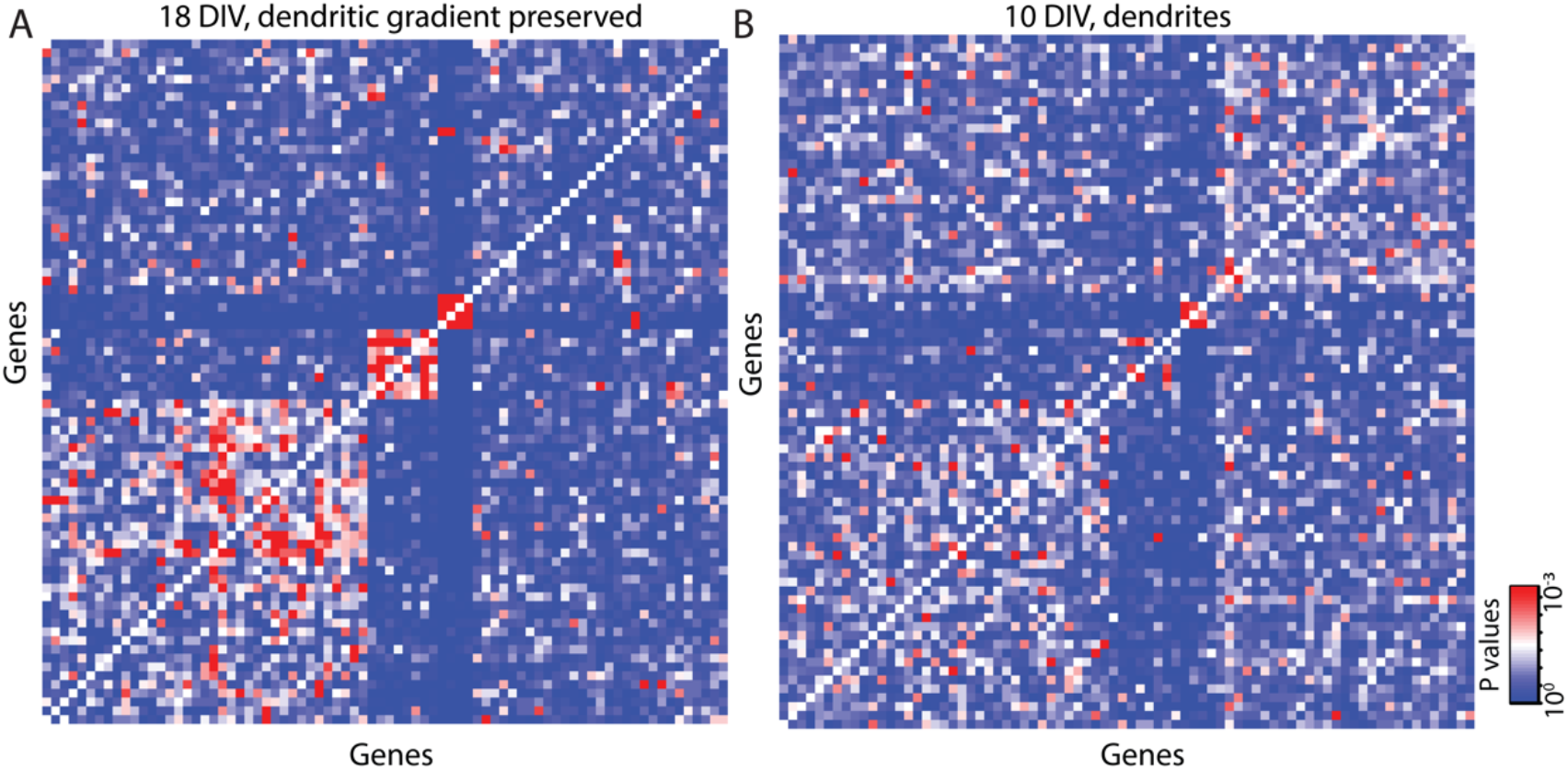
Additional analysis of the spatial association/proximity patterns of RNAs. (**A**) Same as Fig. 4B except that the permutation preserves not only the abundance of each RNA species per dendritic branch but also the gradient along a dendrite for each RNA species. In order to preserve dendritic gradients, dendrites were first segmented into 5-µm-long segments and the permutation was carried out among all RNAs localizing within the same segment. Genes are ordered in the same order as Fig. 4B. (**B**) Same as Fig. 4B but for DIV 10 neurons. Genes are ordered in the same order as Fig. 4B.

